# Bioinformatic pipeline for functional inference of *Rickettsia akari* hypothetical proteins reveals virulence-associated and effector candidates

**DOI:** 10.64898/2026.11.05.764563

**Authors:** Semen Kaliukanov, Frantisek Csicsay, Ludovit Skultety, Marco Quevedo-Diaz

## Abstract

**Background:** *Rickettsia akari*, the causative agent of rickettsialpox, is an obligate intracellular pathogen with a highly reduced genome encoding numerous hypothetical proteins (HPs) of unknown function. Many of these HPs are expressed during host-cell infection, yet their roles in bacterial physiology and potential contribution to virulence remain unclear.

**Methodology/Principal Findings:** We performed an integrative *in silico* characterization of 54 *R. akari* HPs previously identified by mass spectrometry in purified bacterial cells. Using a benchmarked computational workflow, we predicted their physicochemical properties, subcellular localization, putative secretion signals, protein folds, potential interaction partners, and evolutionary conservation. Most HPs were predicted to be thermostable. The majority of HPs were predicted to be cytoplasmic (37.00%) or membrane-associated (29.63%). Predicted features associated with the type IV secretion system were more frequent than those associated with type I secretion. Network and structural analyses identified candidate HPs with features consistent with potential roles in outer membrane assembly, host-cell interaction, and immune modulation, including predicted peptidyl-prolyl cis/trans isomerases, oxidoreductases, methyltransferases, and putative E3 ubiquitin ligases. Several HPs were predicted to be antigenic and showed conservation across multiple rickettsial genomes. Molecular docking and normal mode analysis supported the plausibility of selected predicted protein-protein and protein-ligand interactions. Evolutionary analyses indicated predominant purifying selection among the analyzed HPs. A8GP20 showed elevated sequence divergence and warrants further investigation as a potential effector candidate.

**Conclusions/Significance:** Our findings prioritize A8GPG8, A8GP63, A8GNV4, A8GMM8, A8GM41, A8GMJ7, A8GMI5, A8GP20, A8GN23, A8GMB4, A8GM08, and A8GM39 as promising candidates for experimental validation of their predicted functions and potential roles in *R. akari* biology and pathogenicity. This work provides an integrative and reproducible bioinformatic framework to guide functional discovery in *R. akari* and related neglected intracellular pathogens, and identifies candidate proteins for future studies of rickettsial pathogenesis, antigen discovery, and therapeutic target development.

**Author summary:** Rickettsialpox is a neglected febrile illness caused by the bacterium *Rickettsia akari* and transmitted by mouse mites. Although *R. akari* has a small genome, many of its proteins remain uncharacterized and are annotated as “hypothetical proteins”. These proteins may contribute to the bacterium’s ability to infect host cells and cause disease, but their functions are poorly understood.

We used an integrative computational approach to characterize 54 uncharacterized proteins previously identified in *R. akari* cells by mass spectrometry. Our analyses suggest that many of these proteins are predicted to be stable and associated with the bacterial membrane, whereas several may interact with host cells, potentially through a bacterial secretion system. We identified promising candidates that may contribute to host-cell interactions, immune modulation, outer membrane assembly, and other processes important for bacterial survival. Several proteins were predicted to be antigenic and showed conservation across multiple rickettsial genomes, suggesting potential value for future diagnostic or vaccine studies.

By prioritizing the most promising uncharacterized proteins for experimental validation, our study provides new insights into the biology and pathogenicity of *R. akari* and offers a framework for studying similar proteins in other neglected intracellular pathogens.

## Introduction

*Rickettsia spp*. are Gram-negative, obligate intracellular bacteria transmitted primarily through arthropod vectors [1–3]. Molecular phylogenetic analyses divide the genus into four major groups: the spotted fever group (SFG), the typhus group (TG), the transitional group (TRG), and the ancestral group (AG) [4–7]. Several species within these groups are established human pathogens, including *Rickettsia typhi* (murine typhus), *Rickettsia prowazekii* (epidemic typhus), *Rickettsia conorii* (Mediterranean spotted fever), and *Rickettsia rickettsii* (Rocky Mountain spotted fever) [8–11]. *Rickettsia akari*, another notable human pathogen and member of the transitional group (TRG), causes rickettsialpox and is transmitted by the house mouse mite *Liponyssoides sanguineus*. Unlike most *Rickettsia* species, *R. akari* primarily invades macrophages rather than endothelial cells, leading to fever, headache, and vesicular skin eruptions [8,12–14].

The outcome of rickettsial infection is largely determined by bacterial virulence factors, particularly effector proteins that manipulate host cellular processes to promote invasion, intracellular survival, and dissemination [15–17]. Comparative genomic analyses have indicated that *Rickettsia* species encode core and accessory components of multiple bacterial secretion systems, including the Sec translocon, twin-arginine translocation (Tat) pathway, type I (T1SS), type IV (T4SS), and type V (T5SS) [18]. In addition, a substantial fraction of rickettsial open reading frames encode hypothetical proteins (HPs) of unknown function. Recent evidence has linked HPs in diverse bacteria, including *Rickettsia*, to virulence and pathogenicity [19–22]. Given the highly reduced genome of *Rickettsia*, the conservation of many HPs suggests that they perform essential biological roles [13,23,24]. Because numerous aspects of *Rickettsia*-host interactions remain poorly understood, functional characterization of these HPs is essential [5,25,26].

The obligate intracellular lifestyle of *Rickettsia*, together with limited tools available for genetic manipulation, constrains experimental approaches and increases the value of bioinformatic methods for hypothesis generation. A growing array of computational tools can predict structural and functional features of bacterial proteins, thereby supporting the functional annotation of HPs and informing subsequent experimental work [27–34].

In this study, we characterized the potential functions of 54 HPs in the *R. akari* strain Hartford using an integrative in silico approach. The analyzed HPs were previously detected by mass spectrometry in purified *R. akari* cells [13]. We applied a comprehensive bioinformatic workflow encompassing predictions of physicochemical properties, subcellular localization, conserved domains and motifs, and functional annotations using high-confidence databases and tools. To ensure the robustness of our approach, the analytical workflow was validated against a control set of well-annotated proteins, and only the top-performing resources were retained for final analysis.

We further employed an integrated pipeline that combined fold-type prediction, Gene Ontology categorization, clustering, three-dimensional structural alignment, and conservation and phylogenetic analysis. This approach enabled the prioritization of several HP candidates with features suggestive of potential roles in pathogenicity or other cellular processes. Overall, this study expands the functional annotation of *R. akari* HPs and provides a prioritized set of candidates to guide future experimental validation.

## Materials and methods

### Protein sequence retrieval and database verification

All hypothetical protein (HP) sequences from *R. akari* were retrieved from the UniProt database (http://www.uniprot.org) in FASTA format. Sequence similarity searches were performed using PSI-BLAST against the clustered non-redundant (NR) database, restricted to *Rickettsiales* (taxid:766), and the Protein Data Bank (PDB) database to identify homologous proteins and potential structural templates. Five PSI-BLAST iterations were performed to reduce the likelihood of false-positive identifications.

To assess the predictive performance of the computational tools and databases, a control dataset (n=32) comprising *R. akari* proteins with experimentally supported or well-annotated characteristics was assembled from the Virulence Factor Database (VFDB) (database access: November 2025). Three additional well-characterized proteins, the peptidoglycan-associated lipoprotein, the heat shock protein DnaK, and the chaperonin GroEL, were included for benchmarking (S1 Table).

All the databases and software listed below were accessed between October 2025 and January 2026.

### Characterization of physicochemical properties

Physicochemical parameters for each HP were calculated using the ExPASy ProtParam tool [27], providing theoretical estimates of molecular weight, isoelectric point (pI), aliphatic index, instability index, Grand Average of Hydropathy (GRAVY), and predicted protein half-life. The aliphatic index was used as a sequence-derived indicator associated with thermostability [35], whereas the instability index was used as an indicator of predicted *in vitro* stability [36]. Additionally, the GRAVY index was used to estimate overall sequence hydrophobicity [37].

Protein solubility and expression potential in *Escherichia coli*, a widely used heterologous expression system, were predicted using NetSolP-1.0 [38] and SOLPro [39]. NetSolP-1.0 provides predictions of solubility and expression feasibility, whereas SOLPro provides computational estimates of solubility upon overexpression. Predicted half-lives in mammalian reticulocytes, yeast, and *E. coli* were also obtained.

### Prediction of subcellular localization and secretion pathways

Subcellular localization of the HPs was predicted using a consensus approach based on CELLO, PSLpred, and PSORTb v3.0 [28,33,34]. Membrane topology and orientation were determined using DeepTMHMM and Phobius [40,41], and membrane-protein interactions were modeled using the PPM 2.0 server based on protein surface hydrophobicity [42].

Secretion pathways were inferred using SignalP for Sec/Tat-dependent secretion signals [30] and SecretomeP for potential non-classical secretion [43]. SecretomeP scores were interpreted in the context of predicted localization, membrane topology, and established secretion mechanisms; higher scores indicated a greater likelihood of non-classical secretion.

To assess the performance of localization predictions, receiver operating characteristic (ROC) analysis was performed in GraphPad Prism 5.03 using CELLO and PSLpred outputs. Experimentally validated proteins annotated in UniProt served as controls (S1 Table). PSORTb predictions were evaluated using the same benchmark dataset and cross-validated against UniProt and Argot 2.5 annotations [32]. To integrate localization predictions, the outputs of the aforementioned tools were combined and analyzed using principal component analysis (PCA) and random forest (RF) classification in R v.4.5.1 [44,45]. Finally, the BastionX toolkit [46] was used to assess the effector potential of HPs and to identify candidates associated with rickettsial secretion systems.

### Functional annotation and biological activity prediction

Functional domains and motifs were identified by screening each HP sequence against the HAMAP, CATH, and ScanProsite databases to detect conserved motifs and catalytic signatures [47–49]. Unannotated regions were analyzed using MotifSearch (GenomeNet, Kyoto University; E-value ≤ 0.01) and validated through InterPro [50]. Predicted binding and active sites were identified using the ProFunc server [51].

The functional activity of the HPs was predicted using Argot 2.5 [32] and corroborated by structural and folding analyses. Protein folding patterns were inferred using PFP-FunDSeqE [52]. Argot 2.5 confidence scores >200 and motif alignment E-values <0.01 were considered high-confidence indicators of predicted function.

### Prediction of virulence, antigenicity, and toxicity

The virulence potential of the HPs was assessed using VirulentPred 2.0 and VirulentHunter [29,31]. These predictions were also used to prioritize candidates for further functional and structural analyses, especially when sequence-based functional annotation remained inconclusive.

The antigenic potential of the HPs was estimated using VaxiJen 2.0 with a threshold of 0.4 [53] and further evaluated using AntigenPro [54]. A ROC analysis was performed in GraphPad Prism 5.03 to evaluate the antigenicity predictions. AntigenPro scores were benchmarked against experimentally confirmed antigenic proteins to establish a threshold cut-off that balanced sensitivity and specificity [13,18,55–58].

The potential toxicity of HPs was evaluated using the ToxinPred2 and CSM-Toxin web servers. ToxinPred2, which relies primarily on sequence motifs, amino acid composition, and physicochemical properties, was used to screen for toxin-associated motifs. In parallel, CSM-Toxin, which is based on global sequence relationships and inferred structural features, was used to assess whether the overall properties of the HPs resemble those of harmful proteins [59,60].

### Functional network clustering

Functional relationships among HPs were analyzed by constructing a functional association network using the STRING database (Version 12.0). Topological clustering was performed using the Markov Cluster Algorithm (MCL) integrated within the STRING platform [61]. To evaluate the biological relevance of the resulting clusters, functional enrichment analysis was conducted using STRING-derived network statistics, including enrichment strength (>0.01), connectivity significance (>0.01), and false discovery rate (FDR, ≤0.05) [61]. Clusters that exhibited no significant functional enrichment, sparse connectivity, or consisted exclusively of HPs without known interaction partners were classified as non-informative and excluded from downstream analyses. The remaining clusters were imported into Cytoscape v3.10.4 [62] and Gephi v0.10.1 [63] for visualization and refinement. Molecular Complex Detection (MCODE) with the Haircut algorithm, K-score of 2, node score of 0.2, and degree-based filtering were applied to remove weakly connected nodes. Subnetworks with low node counts, clusters composed exclusively of uncharacterized proteins, or MCODE score < 4.0 were excluded from downstream interpretation.

Several topological parameters were used to characterize the structural and functional properties of the networks [64]. Neighborhood connectivity, defined as the average degree of a node’s neighbors, reflects the connectivity of the local network environment [65–67]. The clustering coefficient quantifies the likelihood that neighboring nodes are connected and reflects network modularity [67,68].

Centrality metrics were used to identify potentially important nodes: (i) betweenness centrality, which measures how often a node lies on shortest paths between other nodes; (ii) closeness centrality, which describes the average shortest path distance from a node to all other reachable nodes; (iii) eigenvector centrality, which assigns importance based on connections to other highly connected nodes; and (iv) authority and hub scores obtained using the HITS algorithm in Gephi v0.10.1, which distinguish nodes that receive many links from nodes that link to many other nodes [65,66,68–70].

### Structural modeling and functional determination

Three-dimensional structures of HPs with the highest MCODE scores were predicted using AlphaFold v6 and evaluated against Protein Data Bank (PDB) templates [51,71,72]. Remote homology modeling and functional inference were performed using Phyre 2.2 [72], with confidence scores >85% considered supportive of the predicted structural assignment. Structural quality was assessed using Ramachandran plots and disorder analysis.

Subsequently, the DALI server was employed to structurally align the predicted HP models against the top Phyre 2.2 hits retrieved from the PDB database. Structural similarity was considered supportive when the DALI Z-score was > 2.0, and the root-mean-square deviation (RMSD) of aligned backbone atoms was < 4.0 Å [73].

### Conservation and phylogenetic analysis

#### Structural conservation analysis

To infer the evolutionary significance of the predicted HP structures, the conservation of structural features and potential ligand-binding sites was analyzed using the ProFunc server. For HPs for which ProFunc was not suitable to identify reliable functional annotations or structural templates, the ConSurf web server was employed. ConSurf calculated position-specific evolutionary conservation scores from multiple sequence alignments, which were then mapped onto the predicted three-dimensional structures to visualize functionally constrained regions [51,74].

#### Sequence conservation and divergence rate assessment

Sequence-level conservation was assessed using MEGA (version 12). All HP nucleotide sequences were searched against the NCBI Non-redundant nucleotide database using tBLASTx, restricted to the order *Rickettsiales* (taxid: 766). Homologous sequences meeting the selection criteria (E-value ≤ 1e^−20^ and identity ≥ 85%) were retrieved. For codon-based analyses, the corresponding coding sequences (CDSs) of the homologs were aligned using the MUSCLE algorithm implemented in MEGA (version 12), maintaining the correct reading frame.

Pairwise evolutionary distances were calculated in MEGA (version 12) using codon-based *p*-distances within the Nei–Gojobori framework [75,76]. The average evolutionary distances were utilized as global indicators of sequence divergence among the identified homologs.

#### Evolutionary rates and selection analysis

Selection analysis was performed using the Nei–Gojobori method to estimate the numbers of synonymous (dS) and non-synonymous (dN) substitutions per site. The dN/dS ratios, Z-scores, and associated *p*-values were calculated using the codon-based Z-test of selection implemented in MEGA (version 12). Positive diversifying selection was evaluated by testing the null hypothesis of neutral evolution (dN = dS) against the alternative hypothesis dN > dS. Purifying selection was evaluated by testing the alternative hypothesis of dN < dS; *p*-values <0.05 were considered statistically significant. Genes exhibiting significant evidence of purifying selection without support for positive selection were classified as evolutionarily constrained [75,76].

Site-specific relative evolutionary rates were estimated using the maximum-likelihood rate estimation procedure in MEGA (version 12) [76–78]which scales rates such that the mean across all sites equals 1.0. To assess evolutionary constraints, the mean and median site-specific rates were recorded, together with the percentage of sites with relative rate ≤ 0.5.

#### Phylogenetic analysis

Phylogenetic trees were constructed using the Neighbor-Joining (NJ) method in MEGA (version 12) [79] and node support was assessed by bootstrap resampling with 1000 replicates [80]. For the genes encoding the HPs, bootstrap support for the node grouping each HP with its closest homolog, along with the corresponding patristic distance, was reported. These metrics summarize the robustness of the inferred relationship and the evolutionary proximity of each focal HP sequence within the phylogenetic topology [76,81].

### Protein structure refinement and validation

Following structural modeling and conservation analyses, molecular interactions between the HPs and their potential functional partners were modeled. The accuracy of the HP tertiary structures was first improved using the GalaxyRefine web server [82]. Refined structures were evaluated using the SAVES v6.0 web server with standard cut-off settings, which assesses stereochemical quality based on Ramachandran plots, sequence-structure compatibility (VERIFY 3D), and nonbonded atomic interaction errors (ERRAT) [83,84]. The overall plausibility of each structure was further assessed using ProSA-web, with a score ≤ −2 serving as a cut-off [85]. From the five refined models generated for each HP, the model with the best validation scores was selected for downstream analyses.

### Functional partner interaction and molecular docking modeling

The refined HP structures were used to model potential molecular interactions. Functional partners and interacting surfaces were selected based on the structural and binding-site analyses conducted in this study. Structural models of interacting partners were obtained from UniProt or the PDB for annotated *R. akari* proteins (GenBank accession: CP000847.1). When no annotated *R. akari* interactor was available, the corresponding homologous protein structure from *E. coli* K-12 (GenBank accession: U00096) was used. A structure of the human mitogen-activated protein kinase 8 (UniProt accession: P45983) was also included as an interactor.

Protein–protein docking was performed using HADDOCK v2.4 software, and protein–ligand docking was carried out using SwissDock with the AutoDock Vina algorithm v3.0 [86–88]. Both tools were used with default docking settings. Docking clusters were evaluated using interaction scores, buried surface area, and interfacial contacts. Protein–protein complexes were further analyzed using the PRODIGY server, which predicts binding affinity based on interfacial contacts [89]. The interactions between amino acid residues of each HP and its predicted partner were explored using PDBePISA [90], providing estimates of interaction free energy, interfacial area, and other parameters relevant to complex formation. Detailed residue-level interactions were visualized using the DIMPLOT and LIGPLOT modules of the LigPlot+ software suite [91]. Docking-derived scores were interpreted as supportive evidence for the plausibility of predicted interactions.

### Normal mode analysis

To evaluate collective motions and structural flexibility of the docked protein complexes, normal mode analysis (NMA) was performed in internal coordinates using the iMOD web server with the Heavy-Atom coarse-grained model as a default parameter [92,93]. The analysis provided estimates of structural mobility, deformability, stiffness, and covariance among residues. Eigenvalues were used to compare the relative stiffness of the predicted complexes, with lower eigenvalues indicating more flexible, low-frequency modes.

iMOD was additionally used to generate feasible transition pathways between selected structural states by iterative deformation along the lowest-frequency normal modes. These analyses were used to assess the conformational plausibility of the modeled complexes and were not interpreted as explicit molecular dynamics simulations.

## Results

### Initial characterization of hypothetical proteins

To evaluate the physicochemical characteristics of the HPs, particularly their predicted solubility and stability, the retrieved sequences were analyzed using the ExPASy ProtParam web server (S2 Table). The results, summarized in Figs 1 and 2, indicated that the analyzed HPs shared several predicted structural features. The predicted *in vivo* half-lives of all HPs in mammalian reticulocytes, yeast, and *E. coli* were 30 h, > 20 h, and > 10 h, respectively. Furthermore, all evaluated proteins were predicted to have high thermal stability; the majority were predicted to be stable *in vitro*, and all displayed hydrophilic sequence features. Regarding solubility, most HPs predicted to be soluble were considered compatible with expression in the *E. coli* system; however, only 26 candidates were predicted to remain soluble upon overexpression.

**Fig 1.**
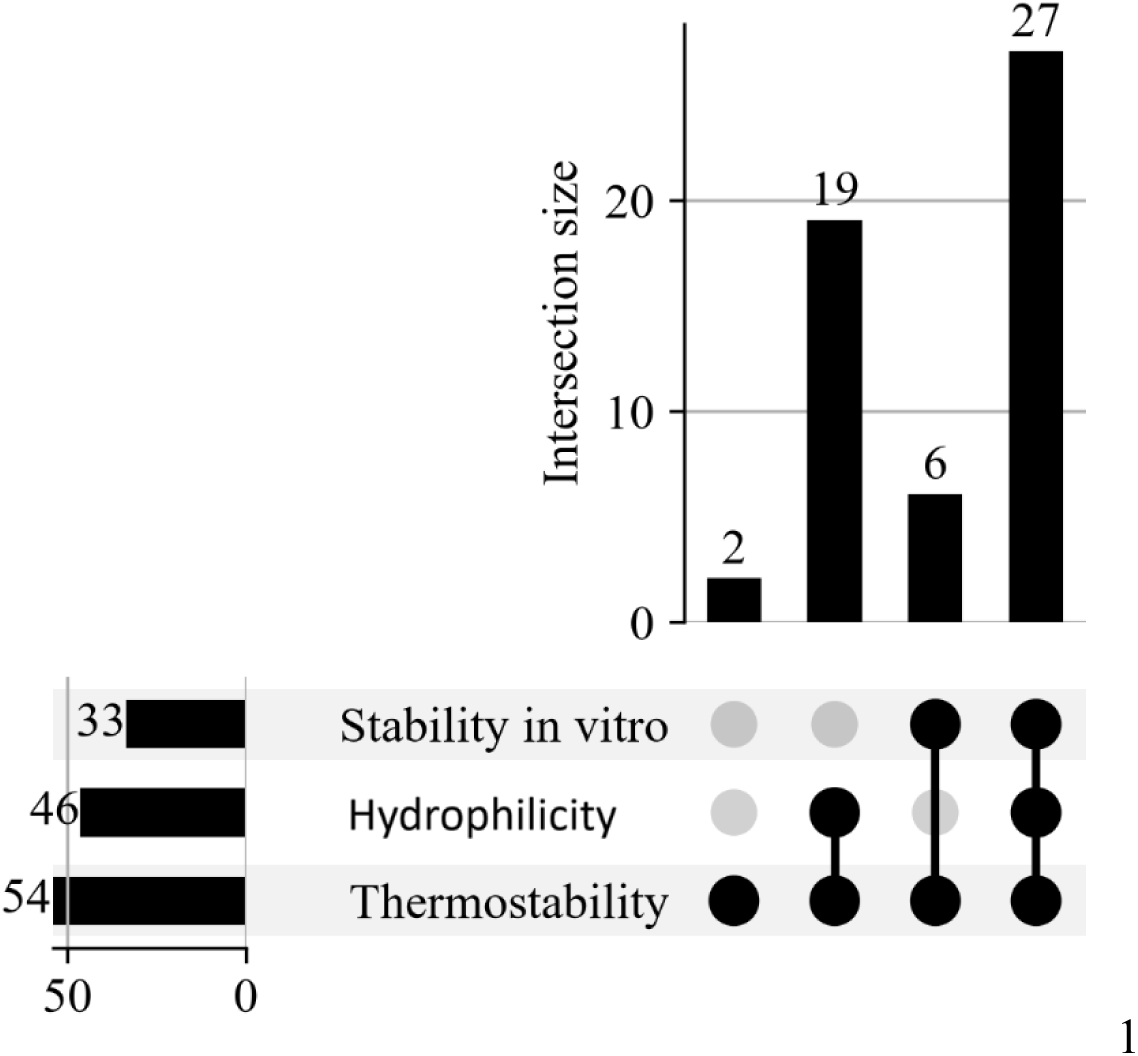
Distribution and co-occurrence of predicted physicochemical properties among *R. akari* HPs.

UpSet plot illustrating the overlap between proteins classified according to predicted thermostability, *in vitro* stability, and hydrophilicity. Horizontal bars represent the total number of HPs assigned to each category, while vertical bars indicate the number of HPs sharing specific combinations of properties. The connected dots below the bars identify the categories included in each intersection.

**Fig 2.**
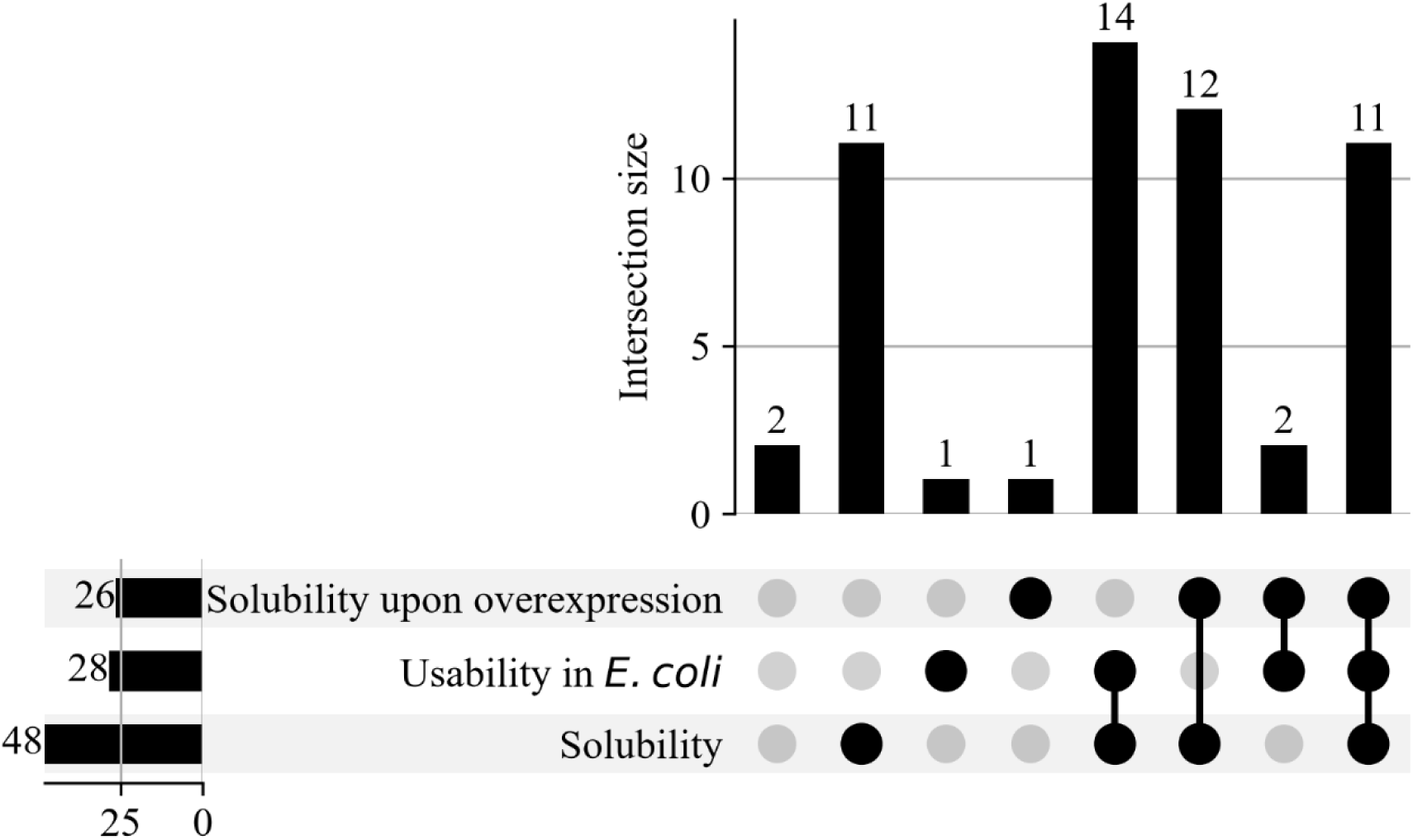
Predicted solubility and recombinant expression characteristics of *R. akari* HPs in *E. coli*.

UpSet plot summarizing the relationships among predicted HPs’ solubility, suitability for heterologous expression, and solubility upon overexpression. The intersection bars indicate the number of HPs meeting each combination of prediction criteria, as shown by the connected dots in the matrix. Horizontal bars show the overall number of proteins assigned to each predicted property.

### Subcellular localization and secretion-associated features

The integrated computational pipeline (described in detail in S1 Appendix) predicted that most HPs were localized to the cytoplasm (37.00%) or associated with the cellular membrane (29.63%), with smaller fractions localized in the periplasm or being surface-exposed. A larger subset of HPs (25.93%) displayed predicted features associated with the T4SS than with the T1SS secretion signals (S3 Table).

Functional predictions identified putative folding patterns, enzymatic activities, and biological roles for the HPs. Several candidates presented features consistent with both predicted virulence-associated functions and metabolic activities. A subset of HPs was predicted to be antigenic based on high scores across multiple immunogenicity and virulence prediction tools (S4 and S5 Tables). Although a large proportion of the HPs were predicted to have virulence-associated characteristics, none exhibited strong predicted toxicity; only minor, unconfirmed indicators of toxicity were identified in isolated cases.

The integrated pipeline highlighted the priority candidates for further investigation. Specifically, the proteins that exhibited combinations of predicted secretion-associated, membrane-associated, antigenic, and virulence-associated features, suggesting that they may warrant experimental evaluation as potential contributors to rickettsial pathogenesis. This portion of the analysis is presented in the S2 Appendix.

### Cluster analysis

Functional protein-protein interaction networks were constructed for each candidate HP using the STRING database to identify HPs with predicted functional associations (S6 Table). To ensure stringent selection, only HPs embedded in dense clusters with significant functional enrichment and high MCODE scores were selected for downstream network topology analysis using Gephi v0.10.1. The resulting topological configurations are compiled in S7 Table and illustrated in Fig 3.

**Fig 3.**
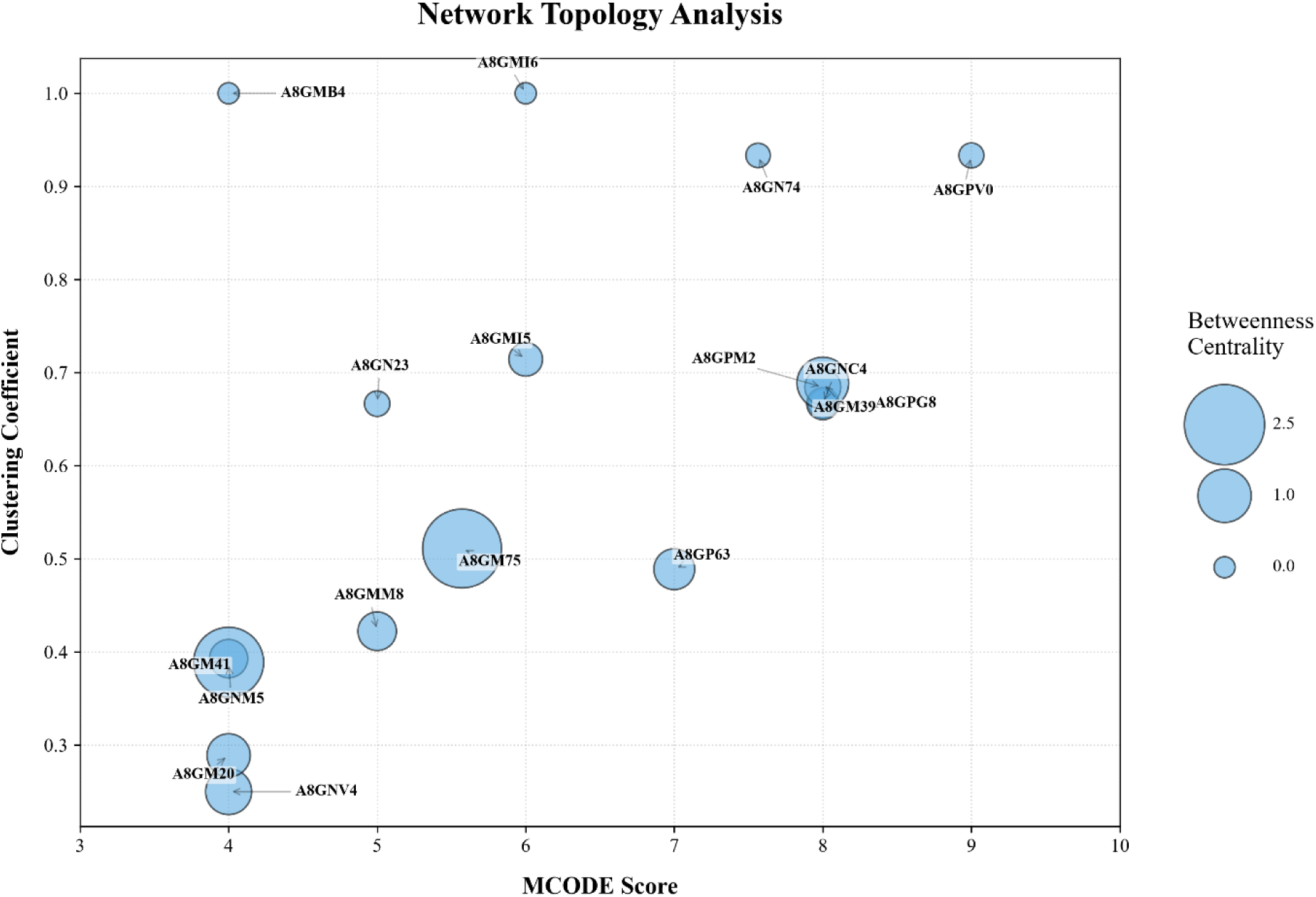
Network topology analysis of the prominent HP clusters identified by MCODE. Each node represents a protein, positioned according to its MCODE score and clustering coefficient. Node size is proportional to betweenness centrality. Proteins with elevated betweenness centrality formed prominent nodes within densely connected clusters, suggesting potential roles as network hubs.

Based on these configurations, distinct predicted functional roles were identified for candidate HPs within their respective networks. A8GNC4, A8GPM2, A8GPG8, A8GN74, A8GM39, and A8GPV0 showed high MCODE scores and clustering coefficients, consistent with potential roles as components of larger functional complexes. In contrast, the high-degree nodes A8GMM8, A8GM75, A8GMI5, and A8GP63 were consistent with potential hub-like roles linking multiple predicted biological pathways. A8GN74, A8GPV0, A8GMB4, A8GMI6, and A8GN23 exhibited high local connectivity but limited global reach, consistent with potential roles in maintaining the structural integrity of specific complexes. A8GNV4, A8GM20, A8GNM5, and A8GM41 occupied more peripheral positions, with low MCODE scores and clustering coefficients, consistent with potential roles as transient linkers or regulatory components rather than structural components.

Modular analysis in Gephi (v0.10.1) grouped these proteins into distinct functional modules, supporting the use of the applied MCODE cut-off to exclude weakly connected nodes (S7 Table). Functional enrichment analysis indicated that the identified clusters were primarily associated with adaptive responses to environmental changes and with predicted regulation of genes encoding cell wall and outer membrane components in Gram-negative bacteria, including predicted *R. akari* orthologs (S7 Table).

### Structural homology analysis

Because three-dimensional structures and active sites are often more conserved than primary sequences, structural homology searches were performed for all HPs embedded within the functional clusters [94–96]. Structurally similar, annotated proteins from diverse bacterial species were identified using the Phyre 2.2 database (S8 Table). In addition, three HPs outside the recognized clusters (A8GP20, A8GMJ7, and A8GM08) were included in the downstream pipeline due to significant and distinctive functional domain predictions (S4 Table). The resulting top matches were evaluated based on confidence scores and sequence identity (S8 Table).

To further assess these initial predictions, three-dimensional structural alignments were performed using the DALI server. A8GPG8, A8GP63, A8GNV4, A8GN23, A8GMM8, A8GMI5, A8GMB4, A8GM41, A8GP20, A8GMJ7, A8GM08, A8GM75, and A8GM39 achieved high scores in both Phyre 2.2 and DALI analyses, supporting their predicted structural configurations (S8 Table and S3 Appendix). Their predicted functions were consistent with the network topology analysis and active-site predictions (E-value ≤ 1 × 10⁻⁵ in ProFunc server), supporting their consideration as candidate network hubs or components of predicted functional complexes (S8 Table). In contrast, Phyre 2.2 models for A8GNC4, A8GPM2, A8GMI6, A8GN74, A8GPV0, A8GNM5, and A8GM20 lacked support from DALI alignments, indicating lower confidence in these specific structural predictions.

Because structural and sequence conservation can mutually support functional predictions, integrated conservation analyses were performed for HPs supported by high-confidence DALI alignments (S8 Table). PSI-BLAST searches indicated that these HPs were conserved across multiple rickettsial genomes, providing additional support for the Phyre 2.2-based structural predictions. A8GP63 and A8GN23 remained annotated as HPs across all examined genomes, possibly reflecting lower structural conservation or divergent evolutionary histories of their predicted three-dimensional structures (S3 Appendix).

### Evolutionary conservation analysis

Consistent with the extensive genome reduction characteristic of rickettsiae, comparative evolutionary analyses indicated that most candidate HPs were conserved (S9 Table) [97]. These proteins exhibited low codon divergence, short phylogenetic distances to their closest homologs, robust bootstrap support, and low mean and median site-specific evolutionary rates; nearly all sites evolved at relative rates ≤0.5. Together, these findings were consistent with functional constraints.

Nei–Gojobori tests supported predominant purifying selection for the examined genes except A8GN23 and A8GP20, which displayed features consistent with relaxed constraint or adaptive evolution. A8GP20 also showed exceptionally high codon divergence and a low number of conserved sites, suggesting rapid evolution or lineage-specific divergence compared with the other HPs analyzed.

### Protein interaction modeling

Molecular docking of HPs was performed using structures refined with GalaxyRefine and supported by structural alignments. As shown in S10 Table and Fig 4, all refined structures had ProSA Z-scores below −4, ERRAT scores generally >90, acceptable Verify3D scores (approximately 60–80% of residues in favorable environments), and Ramachandran plots consistent with acceptable stereochemistry. The HP A8GM75 had a low Verify3D score (10.26%) and was therefore excluded from downstream docking analyses.

**Fig 4.**
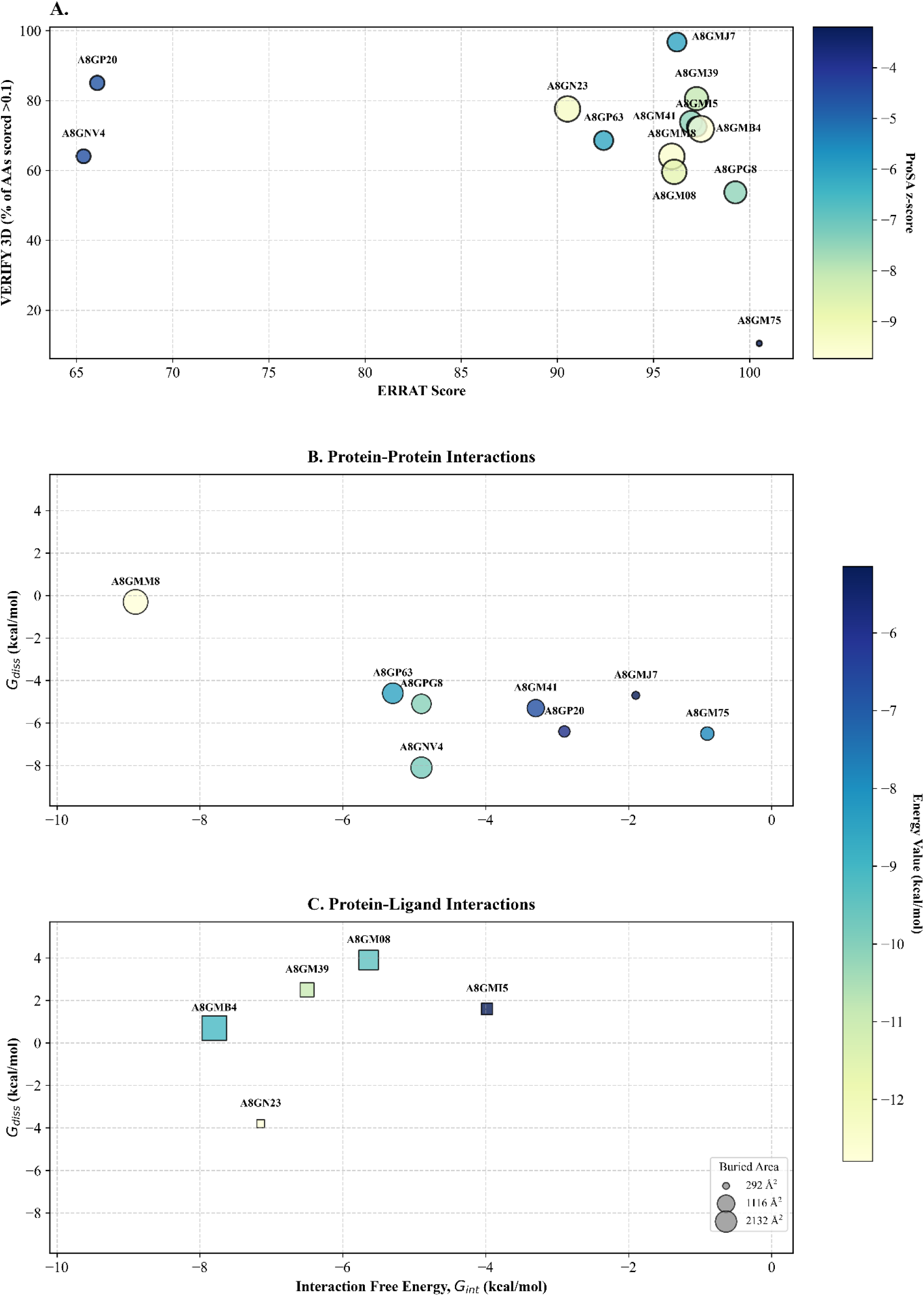
Structural validation and interaction energy landscapes of the refined HPs. (A). Quality assessments of the refined HP structures. The plot assesses the overall stereochemical quality and reliability of the refined HP models by correlating ERRAT scores with Verify 3D percentages of residues with scores ≥ 0.1. The ProSa Z-score is represented by the color gradient bar, which indicates the overall model quality relative to experimental structures. (B) Interaction Energy Landscape Plots combining the numerical features of the selected HPs presented in S10 Table. The graph displays the thermodynamic binding profiles. The axes show the interaction free energy (*G_int_*) in kcal/mol versus the dissociation free energy (*G_diss_*) in kcal/mol. The color bar indicates the energy value (kcal/mol) of the complex as a whole, and the bubble size corresponds to the buried surface area upon complex formation.

Docking cluster analysis generated structural and energetic parameters, including buried surface area, interfacial free energy (ΔG_int_), solvation free energy (ΔG_diss_), and predicted Gibbs free energy (S10 Table). The predicted protein–protein complexes showed ΔG_int_ values ranging from −0.9 to −8.9 kcal/mol and predicted Gibbs free energy ranging from −7.6 to −11.5 kcal/mol, consistent with computationally plausible interactions. Most complexes also exhibited large interaction surfaces and low ΔG_diss_ values (−0.3 to −8.1 kcal/mol), consistent with flexible but plausible associations. A8GMM8 was an exception, displaying a near-zero ΔG_diss_ value, which was consistent with its predicted association with the structurally rigid Bam complex during peptide insertion into the bacterial outer membrane.

Protein–ligand interactions, evaluated with AutoDock Vina, revealed predicted binding energies ranging from −5.1 to −12.8 kcal/mol. Most complexes also showed favorable ΔG_diss_ values (0.7–3.9 kcal/mol), whereas the A8GN23–UDP-GlcNAc complex displayed a lower ΔG_diss_ value (−3.8 kcal/mol). The integrated findings from these computational assessments are summarized in Fig 4.

### Normal mode analysis

To assess the predicted conformational flexibility and collective motions of the docked complexes, normal mode analysis was conducted using the iMOD server. Deformability, eigenvalue, and variance parameters were analyzed (Fig 5) to compare the predicted flexibility and rigidity of the complexes (S10 Table). While deformability analysis predicted regions of increased flexibility, the calculated eigenvalues enabled comparison of the relative stiffnesses of the complexes. Among the analyzed candidates, A8GMJ7 showed the lowest predicted deformability, consistent with its proposed role in iron–sulfur cluster assembly and its predicted interactions with IscS and IscU. Its higher ΔG_int_ and predicted structural rigidity were consistent with the formation of a computationally stable, functionally specialized complex.

**Fig 5.**
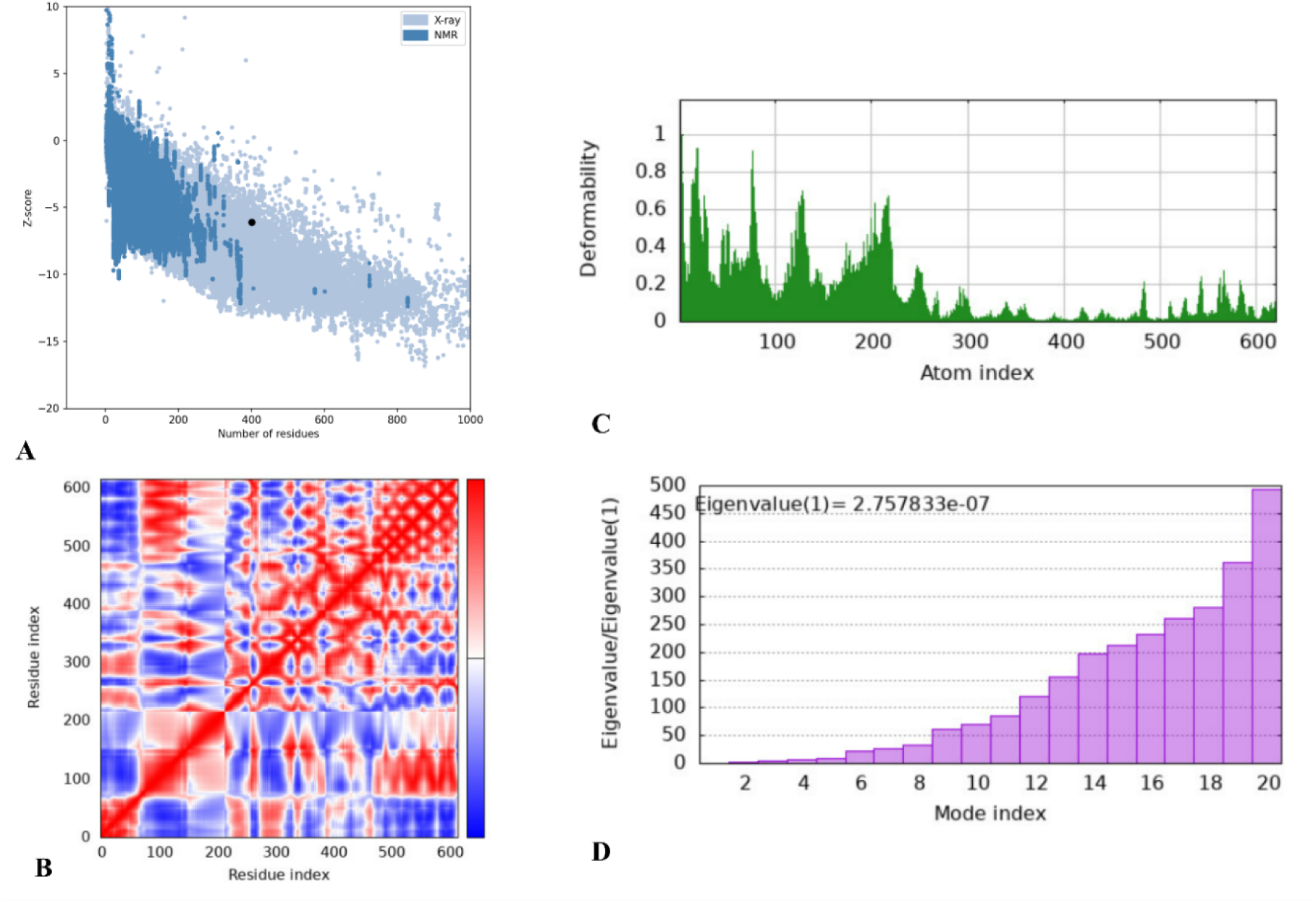
Structural validation and iMODS normal mode analysis of the selected HP. (A) ProSA Z-score plots reflecting overall model quality. (B) Covariance matrix indicating residue fluctuations (red: correlated motion, blue: anti-correlated motion, white: non-correlated motion). (C) Deformability plots. (D) Eigenvalue profiles.

For protein–ligand complexes, low eigenvalues indicated greater predicted flexibility, consistent with potential ligand-binding associated motions. Despite the small molecular sizes of the ligands, the predicted buried surface areas remained substantial. LigPlot+ analysis further identified the predicted noncovalent interaction patterns for each HP–partner pair (S4 Appendix). Collectively, these computational results, although not establishing their biological relevance or binding affinity *in vivo*, supported the structural plausibility of the predicted HP interactions in *R. akari*.

## Discussion

*R. akari* is an obligate intracellular pathogen that predominantly infects macrophages and causes rickettsialpox. The HPs expressed during rickettsial infection represent promising targets for elucidating bacterial virulence mechanisms and life cycle dynamics. In this study, we comprehensively characterized 54 *R. akari* HPs, previously identified by mass spectrometry in purified bacterial cells, using an integrated bioinformatics pipeline to prioritize candidates for downstream experimental validation.

Characterization of the physicochemical properties of these proteins provided useful information for planning downstream experimental assays, including recombinant protein production, purification strategies, and structural or functional analyses (S2 Table and Figs 1 and 2). However, predictions of solubility and compatibility with heterologous expression in *E. coli* are highly sensitive to specific experimental conditions, such as inducer concentration, temperature, and post-induction duration. For instance, although A8GP63 was computationally predicted to be poorly soluble and incompatible with heterologous expression, it was successfully expressed and purified in high yield from *E. coli* BL21(DE3) under optimized conditions [13]. This example highlights the need to interpret solubility predictions as guides for experimental design rather than as definitive predictors of protein behavior.

The majority of the examined HPs were predicted to have high thermostability and *in vitro* stability, suggesting potential structural robustness in biochemical assays. Most candidate proteins displayed negative GRAVY values, which are generally associated with a more hydrophilic overall sequence character. When considered together with predicted transmembrane regions and membrane-topology features, these results suggest that a subset of the HPs may be associated with cellular membranes or periplasmic compartments.

Given the degree of subcellular organization in bacteria, prediction of subcellular localization can help generate hypotheses about protein function [98–101]. Most evaluated HPs were predicted to be cytoplasmic and globular; however, the identification of transmembrane regions or non-cytoplasmic membrane topologies supported potential periplasmic or membrane-associated localization for a subset of proteins (S3 Table).

Bacterial secretion is an important feature of many effector proteins, although the presence of a secretion-associated signal alone does not demonstrate effector activity. We therefore integrated SignalP and SecretomeP predictions with membrane topology, orientation, and localization consensus data derived from CELLO, PSORTb, and PSLpred [30,102,103]. This integrated approach was further supported by PCA and RF modeling. When combined with BastionX screenings, HPs with moderate or elevated T4SS-associated scores, predicted secretion signals, and non-cytoplasmic localization were prioritized as candidate effectors. Notably, several HPs with only moderate or weak T4SS-associated scores also displayed these structural features, suggesting that reliance on rigid algorithmic cutoffs may exclude biologically relevant effector candidates.

Several T4SS candidates identified in this study (A8GNV4, A8GNB3, A8GP22, A8GN63, A8GNW9, A8GMW9, A8GPI5, A8GPV0, A8GM39, A8GN74, A8GNX2, A8GMI6, A8GMB4, and A8GN23) were previously classified as likely *R. akari* effectors using the OPT4e framework and possess established orthologs in *Rickettsiales* [22]. Within this cohort, A8GNB3 has been linked to pathogen-specific effector functions, whereas A8GNW9, A8GN63, A8GMW9, A8GNV4, A8GM39, A8GN74, A8GPV0, and A8GMI6 were previously predicted to be core T4SS effectors. Although minor discrepancies exist among computational approaches, the overlap between our predictions and previous analyses supports prioritizing these proteins for future functional characterization.

Before reaching their final subcellular destinations, proteins must adopt specific three-dimensional folds that are intrinsically linked to their operational mechanisms [104,105]. Although protein folds can provide useful clues to molecular function, structural similarity alone does not definitively establish function. The most prevalent structural configuration among the analyzed HPs was the Immunoglobulin-like (Ig-like) fold, which is found in bacterial adhesins and outer-membrane proteins and has been associated with host-cell interactions [106,107]. This was followed by the oligonucleotide/oligosaccharide-binding (OB) fold, typically associated with nucleic acid or carbohydrate binding and host-cell signaling cascades [108,109]. Other frequently observed architectures included the four-helical cytokine fold, DNA-binding three-helical bundle, and four-helical up-and-down bundle, which have been associated with signal transduction, DNA metabolism, or membrane-associated processes [110–113].

Several identified folds were consistent with the predicted functional annotations. For example, an EF-hand motif was recognized in a putative iron-sulfur cluster protein, whereas a conA-like lectin/glucanase fold was identified in a predicted oxidoreductase, an assignment supported by both CATH and InterPro databases. In addition, a triosephosphate isomerase (TIM)-barrel fold was identified in A8GM08, consistent with its predicted dGTPase activity [114–117].

A substantial portion of the HPs were predicted to possess acyl-transferase activity. This broad functional category is implicated in diverse biological pathways, complicating precise functional assignments (S4 Table) [115,118,119]. Nevertheless, several predicted activities could be relevant to bacterial physiology or pathogenesis. Predicted peptidyl-prolyl cis/trans isomerases may contribute to outer-membrane protein folding and assembly through interactions with DnaK, DegP, and the BAM complex, and disruption of related proteins has been associated with impaired virulence-associated phenotypes [120–122]. Predicted oxidoreductases may contribute to disulfide bond formation required for the maturation of secreted proteins and pili [123–125], whereas proteins involved in heterodimerization may participate in processes relevant to the assembly or activity of bacterial toxins[126–129]. These functional assignments remain hypothetical and require experimental validation.

Network topology analysis further refined the initial functional annotations. The predicted functional assignments for A8GPG8, A8GP20, A8GP63, A8GNV4, A8GN23, A8GMM8, A8GMI5, A8GMB4, A8GM41, A8GMJ7, A8GM39, and A8GM08 were substantiated by complementary structural, domain-based, and network-based evidence. Molecular docking provided additional support for the structural plausibility of selected predicted interactions. The functional clusters containing these HPs were associated with outer membrane assembly, bacterial physiological adaptation, and host-pathogen interactions. Six proteins (A8GP20, A8GMB4, A8GN23, A8GP63, A8GM39, and A8GNV4) were linked to predicted virulence-related activities (S5 Table), whereas the remaining HPs were predominantly associated with core metabolic processes (S4 Table).

The biological relevance of these HPs is further supported by conservation analyses. Although structural and binding-site conservation varied across the dataset, PSI-BLAST queries indicated broad conservation within rickettsial genomes. Moreover, comparative evolutionary analyses also provided evidence that the majority of these proteins evolve under purifying selection. Taken together with predicted virulence-associated features and structural similarity to annotated bacterial proteins, these candidates represent high-priority targets for experimental functional characterization.

A8GP20 emerged as an evolutionary outlier with substantial functional ambiguity (S9 Table). Phyre 2.2 modeling indicated structural homology to AlbG, suggesting a possible role in DNA protection, whereas functional domain analyses were consistent with SopA-like E3 ubiquitin ligases, a signature associated with some virulence effectors (S4 Table). The presence of a Sec/SP2 signal peptide and a lipoprotein attachment site further pointed to potential membrane targeting and surface exposure. This combination of features is consistent with a possible moonlighting or context-dependent function, although this interpretation requires experimental testing [130–132].

This integrative *in silico* analysis has several limitations. Structural and fold predictions rely on computational models and may not capture dynamic properties, such as conformational flexibility, ligand-induced allosteric changes, or protein interactions within the intracellular environment of *R. akari*. Functional annotations based on remote homology, domain prediction, network analysis, and docking remain hypothetical and may be affected by incomplete databases, limited sequence diversity, or inaccurate structural models. In addition, secretion and effector predictions cannot demonstrate protein translocation or host-cell interaction. Therefore, the predicted properties of these HPs should be interpreted as a structured set of testable hypotheses designed to guide and prioritize future experimental validation.

## Conclusions

This study establishes an integrative *in silico* framework to characterize 54 HPs previously identified by proteomic analysis of purified whole-cell *R. akari*. By combining physicochemical, subcellular localization, secretion-associated, structural, interaction-network, and evolutionary analyses, we generated putative functional annotations and prioritized candidate proteins for experimental validation. Several HPs were predicted to be stable, membrane-associated, or potentially involved in host-cell interaction and secretion-related processes. A subset of candidates was associated with functional interaction networks that are potentially relevant to bacterial adaptation and pathogenicity, warranting further investigation as virulence-associated or effector proteins.

The integration of subcellular localization, secretion prediction, and analyses of T4SS- and T1SS-associated proteins enabled targeted prioritization of candidate effector proteins. This approach highlights proteins supported by multiple complementary bioinformatic features, including candidates with moderate individual predictions that might otherwise be overlooked. Our findings prioritize A8GPG8, A8GP63, A8GNV4, A8GMM8, A8GM41, A8GMJ7, A8GMI5, A8GP20, A8GN23, A8GMB4, A8GM08, and A8GM39 as promising candidates for experimental validation of their predicted functions and potential roles in *R. akari* biology and pathogenicity. This work provides a structured bioinformatic framework to guide functional discovery in *R. akari* and related neglected intracellular pathogens and identifies candidate proteins for future studies of rickettsial pathogenesis, antigen discovery, and therapeutic target development.

## Data availability statement

All relevant data are included within the paper and supporting files.

